# Agriculture alters protein evolution of nitrogen cycling genes in soil bacteria at a global scale

**DOI:** 10.1101/2025.05.07.652758

**Authors:** Timothy M. Ghaly, Bhumika S. Shah, Nicholas V. Coleman, Liam D. H. Elbourne, Johannes J. Le Roux, Michael R. Gillings, Ian T. Paulsen, Sasha G. Tetu

## Abstract

Humans are the world’s greatest evolutionary force. Yet, our impacts on the evolution of Earth’s microbiomes and their biogeochemical processes remain poorly understood. Notably, the overlooked potential for the intensive use of agricultural fertiliser to drive evolutionary changes in soil nutrient cycling genes warrants urgent attention. Here, analysing >2,500 soil metagenomes from across the globe, we identify increased rates of diversifying positive selection on genes involved in the reduction of nitrate (a key component of nitrogen fertilisers) in agricultural, but not natural land systems. Altered selection on genes encoding the respiratory nitrate reductase were specific to Burkholderiales, a major group of denitrifying bacteria. We provide evidence that agriculture is driving evolution of protein regions implicated in substrate access to the enzyme’s active site, possibly resulting in increased rates of nitrate reduction. We hypothesise that increasing substrate turnover would be evolutionarily advantageous under excess nitrate availability, ultimately enhancing growth rates despite potential enzymatic trade-offs. As Burkholderiales are dominant denitrifiers globally, such evolutionary consequences of agriculture on this lineage could have cascading ecological impacts. These findings indicate that anthropogenic selection can alter protein-level evolution of vital microbial biogeochemical processes.

## Introduction

In the face of global environmental change and increasing food demand [1], there is an urgent need to maintain food production within the safe operating confines of planetary boundaries [2-6]. The nitrogen (N) and phosphorus (P) cycles represent pivotal boundaries within this context. Transgression of N and P boundaries can substantially alter Earth system health and resilience [7]. Severe disruptions to N and P flows, which are generally localised to intensive agricultural zones, can extend to impact nutrient cycles on a global scale [7]. Humans now input 110 teragrams (Tg) of N per year from synthetic fertilisers alone, with an upward trajectory in global usage [8]. This places humanity in the high-risk zone of the N planetary boundary, set at a total input of 134 Tg N·year^−1^ [5]. Concerningly, P input through fertilisers is now 14.2 Tg P·year^−1^ [9, 10], which already exceeds the P planetary boundary of 6.2 – 11.2□Tg P·year^−1^ [7, 11].

Despite known ecological impacts, possible evolutionary consequences of agrochemical use on microbial biogeochemistry have not been fully explored. Humans are the most powerful evolutionary force on the planet [12], with pollution being a major evolutionary driver [13]. In particular, intensive agricultural inputs of N and P compounds, which flood agroecosystems with the substrates and products of microbial nutrient cycling pathways, could be exerting strong selection on nutrient cycling genes. Microbial biogeochemistry, mediated by metabolic oxidation-reduction (redox) reactions, drives nutrient fluxes that shape the dynamics of entire biomes. Within terrestrial systems, this metabolic activity is the foundation of ecosystem functioning, and is tightly linked with aboveground productivity and biodiversity [14].

Important, yet unaddressed, questions surround the evolutionary consequences of anthropogenic activities altering the dynamics of redox reactions that have been key steps in microbial metabolism for billions of years. Changes to selection patterns on important nutrient cycling enzymes could have substantial environmental and ecological impacts. Key insights have been made regarding anthropogenic impacts on the composition of nutrient cycling microorganisms and enzymes in soils globally [15-18]. However, the potential evolutionary impacts of such global change factors have remained unexplored. Determining how selection regimes on N and P cycling genes differ between agricultural and natural land systems, and between microbial taxa, is essential to better understand, and therefore manage, N and P pollution.

Here, we used a global dataset of more than 2,500 soil metagenomes to examine agriculture-specific signals of selection for N and P cycling genes. We found that genes encoding the respiratory nitrate reductase (Nar) displayed higher rates of diversifying positive selection in agricultural versus natural land systems. This altered selection is specific to Burkholderiales, a globally distributed bacterial taxon associated with plant and soil health [19-22], and a major driver of denitrification [23-27]. We provide evidence that agriculture is driving the evolution of Nar enzyme-substrate dynamics, likely leading to increased rates of nitrate reduction. Together, our findings provide the first evidence of agricultural practices influencing the evolution of microbial genes globally, with considerable environmental and ecological implications.

## Results and Discussion

Here, we analysed a global dataset of 2,545 soil metagenomes sourced from agricultural, grassland, temperate forest, and tropical forest land systems (Fig. 1a; Supplementary Table S1), to examine agriculture-specific signals of adaptive selection for N and P cycling genes. To infer patterns of selection (i.e., positive versus purifying selection), we estimated the ratio of amino acid-replacing non-synonymous (d_N_) to silent synonymous (d_S_) mutations on a per codon basis (Fig. 1b). A d_N_/d_S_ ratio (hereafter denoted as ω) greater than one, indicates positive (diversifying) selection (i.e., selection to change the amino acid at that site), while ω < 1, indicates purifying selection (i.e., selection to maintain the amino acid). Using this approach, we estimated codon-wise ω values for 20 nitrogen and 18 phosphorus cycling genes (Fig. 1c; Supplementary Table S3).

**Fig. 1.**
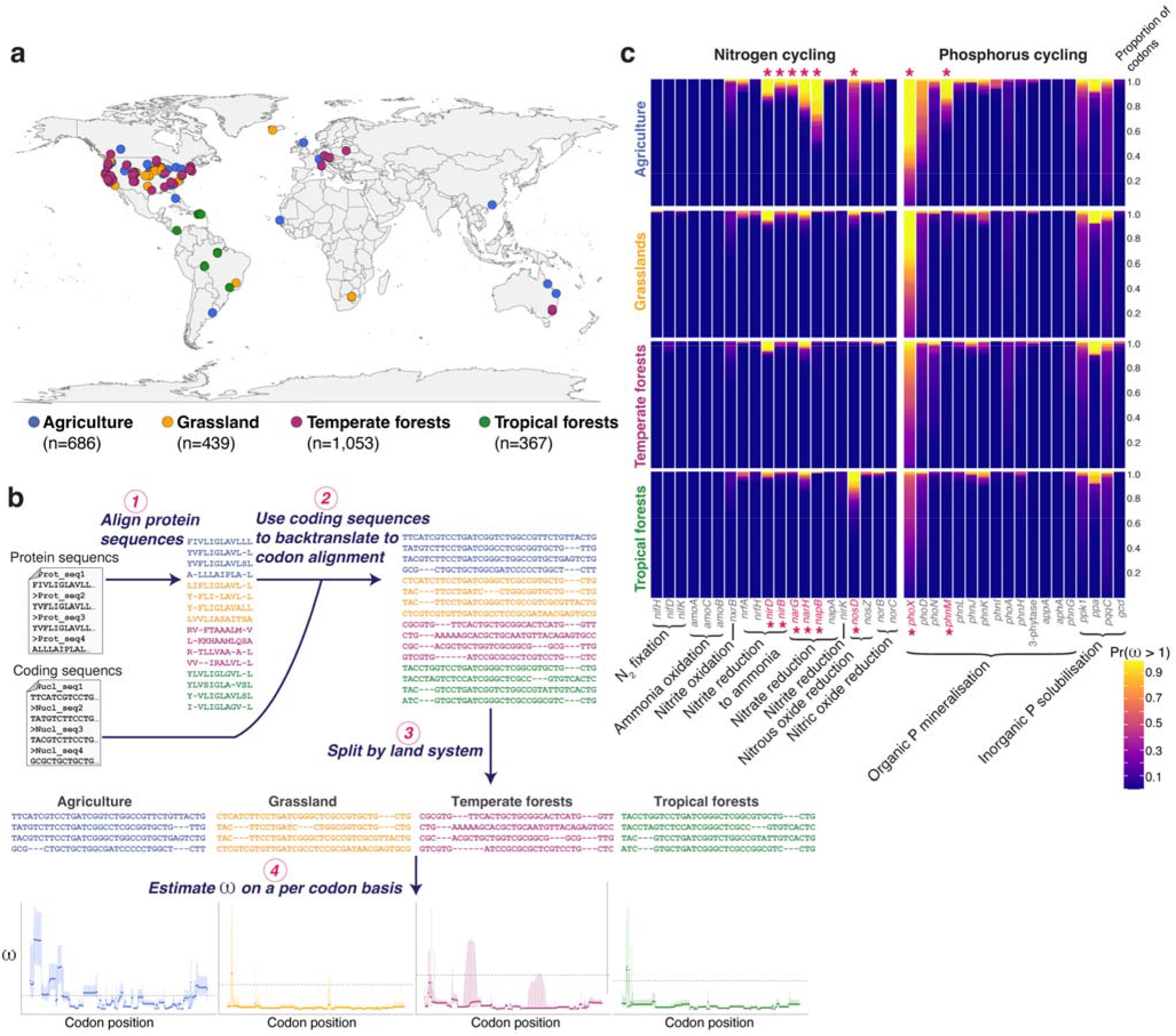
Estimating ω on a per codon basis for soil nitrogen (N) and phosphorus (P) cycling genes across land systems. **a**, Global distribution of bulk soil metagenomes used for screening N and P cycling genes. Sample locations are coloured by land system, with the number of metagenomes per land system displayed in parentheses. **b**, Analysis workflow for estimating ω, which involved: *1 -* alignment of all protein sequences for a given gene, *2* - ‘backtranslation’ to a codon-based alignment using the corresponding coding sequence for each protein, thus ensuring no gaps within any codon, *3* - separating aligned sequences based on their respective land systems, and *4* - estimating ω on a per codon basis for each land system separately (see Methods for more details). This generated, for each codon, a ω point estimate (points), and a 95% credibility interval (error bars) **c**, Stacked bar charts displaying the proportion of codons for each gene with a given strength of evidence for positive selection, defined as the posterior probability (Pr) of ω > 1 for a given codon. Codons with a low probability of positive selection are coloured blue, while those with a high probability of positive selection are yellow. Genes that differed significantly in the proportion of codons under positive selection (defined as a 95% credibility interval for ω > 1) between land systems are displayed in pink text with asterisks (pairwise χ^2^ tests, Benjamini-Hochberg-adjusted p-values < 0.01).

### Land system- and taxon-specific positive selection on respiratory nitrate reductase

Eight N and P cycling genes (*narG, narH, napB, nirB, nirD, nosD, phnM, phoX*) had a significantly greater proportion of codons under positive selection (ω > 1) in at least one land system (Fig. 1c; Supplementary Fig. S1; χ^2^ tests, Benjamini-Hochberg-adjusted p-values < 0.01). The remaining genes displayed similar patterns of purifying selection (ω < 1) across all land systems (Fig. 1c), and almost all had near identical codon-wise ω values across land systems (Supplementary Figs. S2-S3).

Next, we explored the role of taxonomy on the observed land system-specific selection for these eight genes. Codon-wise ω values were re-calculated for the eight significant genes within each land system, restricting analyses to individual taxa (see Methods for details). We searched for genes within taxonomic lineages that exhibited signals of positive selection in any one land system (Supplementary Fig. S4). This revealed two genes, *narG* and *narH*, within Burkholderiales, that had significantly greater proportions of codons under positive selection in agricultural, but not natural land systems (χ^2^ tests with Benjamini-Hochberg-adjusted p-values < 0.0095; Figs. 2b,c). Burkholderiales were one of the four major taxa (together with Mycobacteriales, Rhizobiales and Enterobacteriales) identified to be *narG* and *narH* carriers (Fig. 2a). To further validate these findings, we pooled all Burkholderiales sequences from samples belonging to the three natural land systems (i.e., grassland, temperate forests, and tropical forests) for *narG* and *narH*, and reperformed the ω analysis. This was to maximise Burkholderiales *narG* and *narH* codon sequence variation from non-agricultural sites, and thus enhance the statistical power to detect codons under positive selection in non-agricultural soils [28]. Despite this, however, we found no increase in the number of codons under positive selection in non-agricultural soils, and still observed a significant increase in the proportions of codons under positive selection in agricultural compared to the other land systems combined (χ^2^ tests, Benjamini-Hochberg-adjusted p-values ≤ 0.01; Supplementary Fig. S5).

**Fig. 2.**
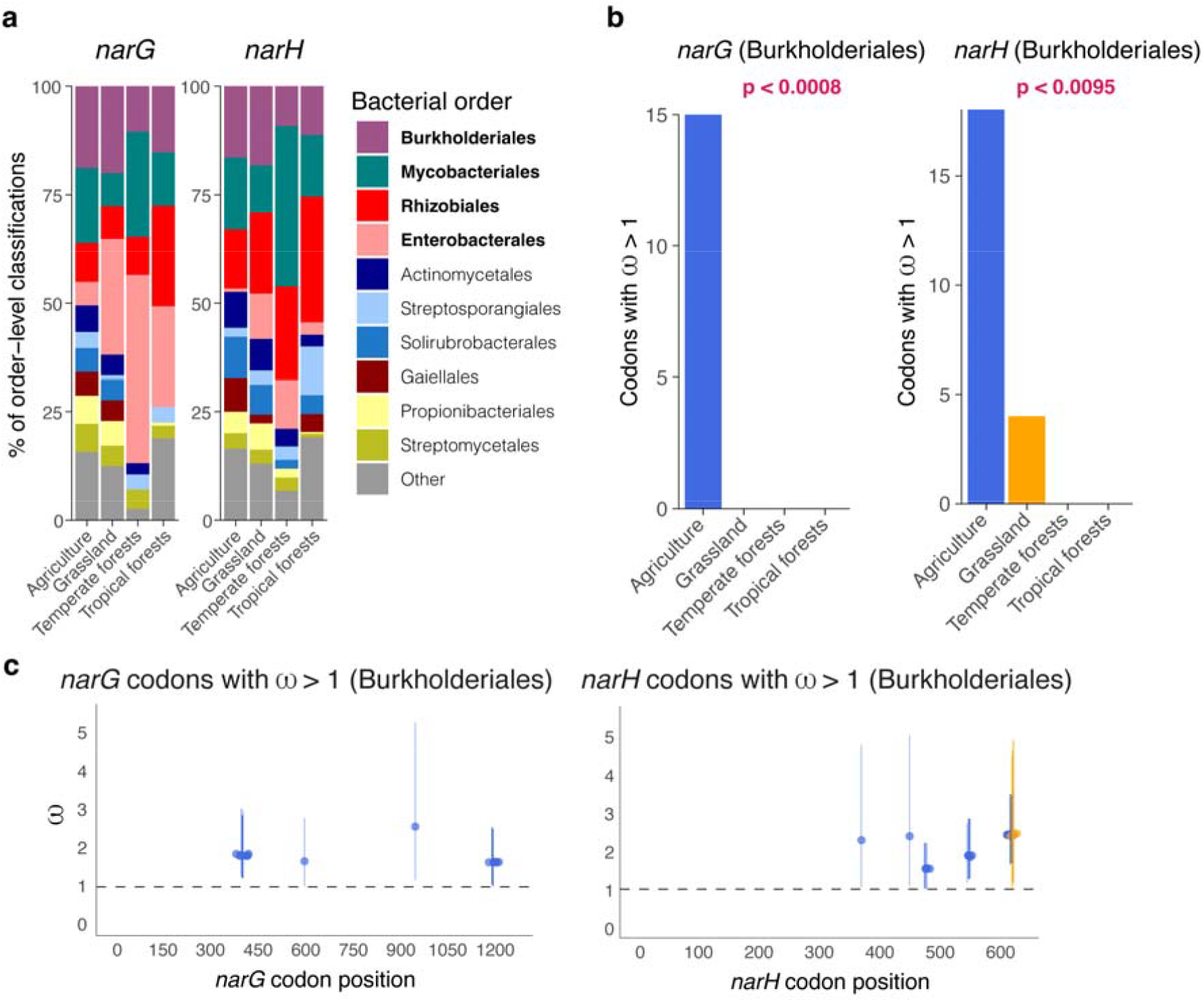
Land system- and taxon-specific positive selection on Nar respiratory nitrate reductase genes. **a**, Order-level profiles of taxa encoding *narG* and *narH* among each land system. The four most prevalent taxa (labelled in bold font) were used for taxon-specific ω analyses (See Supplementary Fig. S4). **b**, The number of *narG* and *narH* codons under positive selection for Burkholderiales within different land system. There were significantly more *narG* and *narH* codons under positive selection in agricultural soils compared to all other land systems (Pairwise χ^2^ tests with Benjamini-Hochberg p-value correction). **c**, All Burkholderiales *narG* and *narH* codons under positive selection along the length of the genes, coloured by land system, with points representing posterior median ω values, and bars represent their corresponding 95% Bayesian credibility intervals. Points are jittered for clarity.

The genes *narG* and *narH* encode subunits of the respiratory nitrate reductase (Nar), which drives anaerobic reduction of nitrate to nitrite. Nitrate is a key component of N fertilisers, and thus, the flooding of substrate in agroecosystems could be driving the evolution of specific protein residues. The increased rate of Nar positive selection in agricultural soils was observed only for Burkholderiales (Fig. 2b-c), which was one of the major taxa contributing to the pool of *narG* and *narH* sequences across land systems (Fig. 2a). Burkholderiales are ubiquitous in soils globally, are key players in Earth’s nutrient cycles, and are intimately linked to plant and soil health [19-27].

### Burkholderiales NarG residues under selective change are associated with the substrate channel

To better understand the contribution to Nar protein function that the sites under positive selection might have, we mapped their locations to the corresponding crystal structures of NarG (PDB accession 1R27_A) and NarH (1R27_B) from *Escherichia coli*. For this, we examined the location of all residues that had a posterior probability of positive selection greater than 90% (i.e., Pr(ω>1) > 0.9).

NarG residues undergoing selective change in agroecosystems suggest possible adaptation for altered Nar enzyme kinetics (Fig. 3). The solved crystal structure of NarG, which is the catalytic subunit of Nar, reveals a narrow substrate channel leading down to the active site [29, 30]. We found that the codons corresponding to active site residues are under strong purifying selection (ω ≈ 0; Fig. 3a). In contrast, those under positive selection, which are spread along the length of the *narG* sequence (Fig. 3a), encode residues that fold together to form two clusters that flank the entrance to the substrate channel (Fig. 3b). This indicates that evolution at these residues is likely impacting enzyme-substrate interactions. Another cluster of NarG residues under positive selection is in a region involved in docking with NarH, the Nar electron transfer subunit (Fig. 3b). However, these residues are not in close proximity to the interaction interface between NarG and NarH subunits (Supplementary Fig. S6), making the functional significance of these residue changes less clear. Likewise, the NarH residues under positive selection do not appear to be associated with NarG-NarH subunit interactions, or its electron transfer pathway (Supplementary Fig. S6).

**Fig. 3.**
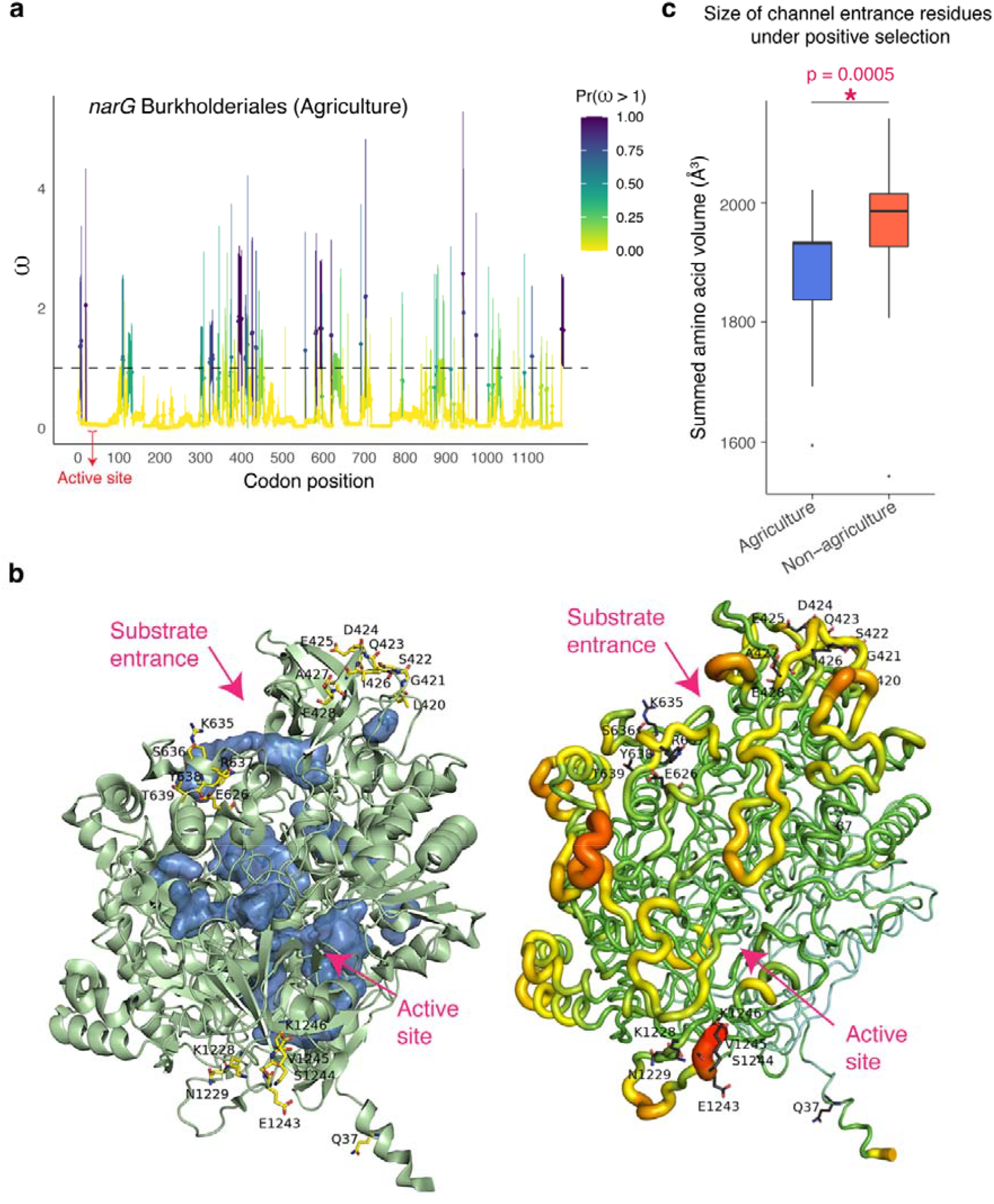
Burkholderiales NarG sites under positive selection in agricultural soils. **a**, posterior median ω values per codon (points), with corresponding 95% credibility intervals (error bars) along the length of Burkholderiales NarG sequences from agricultural soils. Colour scale bar indicates the posterior probability (Pr) that ω is greater than one at that codon (i.e., Pr(ω > 1)). Codons corresponding to active site residues are displayed by the red arrow. **b**, The left structure depicts the solved NarG crystal structure from *Escherichia coli* (PDB accession 1R27_A) as a green ribbon diagram. Sites under positive selection (i.e., Pr(ω > 1) > 0.9) are shown as yellow sticks, and the substrate channel cavity is filled in blue. Pink arrows indicate the entrance to the substrate channel and the approximate location of the active site. The structure on the right displays a B-factor putty representation of NarG (1R27_A), illustrating dynamic mobility of the protein. Orange to red colours, and a wider tube, signify regions with higher B-factors, and thus greater flexibility, whereas shades of blue, and a narrower tube, indicate regions with lower B-factors, and are thus more rigid. Note that the sites under positive selection are on highly flexible regions of the protein. **c**, Boxplots comparing the total size (left boxplot) and charge (right boxplot) of all residues flanking the entrance of the substrate channel that are under positive selection. Total size is displayed as the summed amino acid volume (Å^3^). Note, this region is significantly smaller, and contains significantly fewer negatively charged amino acids, in NarG proteins from agricultural soils than those from non-agricultural soils (Wilcoxon rank sum tests with continuity correction, p ≤ 0.001).

Notably, the NarG sites under positive selection are in highly flexible regions of the protein (Fig. 3b), suggesting that those at the channel opening play a role in structural plasticity required for substrate access to the active site. This further suggests that agriculture-specific selection is likely impacting enzyme-substrate dynamics.

To gain further insights, we next examined the specific changes in amino acids and the physicochemical properties of the NarG residues flanking the entrance to the substrate channel. We investigated amino acid size (defined as its volume in Å^3^), charge, and hydrophobicity. We found that of these properties, amino acid size appears to be under selective change, being overall smaller in NarG proteins in agricultural soils compared to non-agricultural soils, with a significant reduction in total amino acid volume (Fig. 3c; Wilcoxon rank sum test, p=0.0005). No difference was detected for hydrophobicity (Wilcoxon rank sum test, p=0.1132), and on average, there was only one less negatively charged residue (out of 15) in agricultural NarG proteins. This indicates that selective pressure in agricultural soils favours the substitution of larger amino acids at the channel entrance with smaller counterparts, resulting in the widening of the Burkholderiales nitrate reductase substrate channel.

### Hypothesis of agriculture-specific selection for altered Nar enzyme kinetics and increased substrate turnover

We hypothesise that the amino acid changes described above could drive faster rates of nitrate reduction since a larger channel opening is associated with a faster turnover of substrate [31]. We propose that this would be evolutionarily advantageous in agricultural soils with sustained excess nitrate availability (Fig. 4) [32]. Here, the reduction of more nitrate molecules per unit of time would result in a faster respiration rate and, subsequently, faster growth rate. It is worth noting, however, that such an increase in enzymatic reaction rate often comes with a trade-off, negatively impacting substrate affinity (Fig. 4a) [33]. Higher substrate affinity allows an enzyme to efficiently bind and catalyse its substrate even at low concentrations [33]. When substrate concentrations are high, affinity is less important, and thus, the favouring of substrate turnover becomes a more effective strategy. Therefore, in environments where resources are abundant, such as nitrate in agricultural soils [32], there is likely a selective advantage to increase substrate turnover, even if this comes at the cost of substrate affinity, ultimately maximising growth rate (Fig. 4b).

**Fig. 4.**
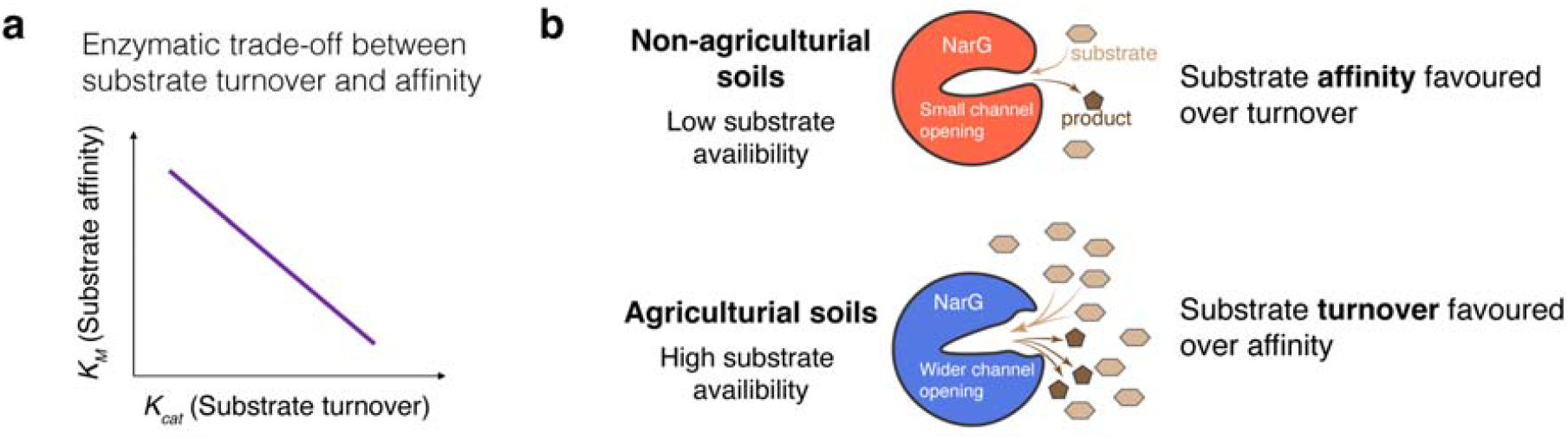
Hypothesis of agriculture-specific selection for altered Nar enzyme kinetics and increased substrate turnover. **a**, Enzymatic trade-off between substrate turnover and affinity – increased substrate turnover generally comes at the cost of reduced substrate affinity. **b**, Proposed hypothesis for NarG positive selection in agricultural soils. When substrate availability is low, then substrate affinity will likely be favoured, allowing the enzyme to bind and process what little substrate there is more effectively. However, when substrate availability is high, then affinity might be less important than substrate turnover, favouring the reduction of more nitrate molecules per unit of time, resulting in a faster respiration rate and, subsequently, faster growth rate.

This carries particular weight for Burkholderiales, a major driver of denitrification in many environments, including soils [23-27], which comprises a step-wise series of reductive reactions beginning with the reduction of nitrate [34]. Nitrate reduction is also the first step in the dissimilatory nitrate reduction to ammonium (DNRA) pathway [34]. Both processes compete for nitrate. However, the energy yield from DNRA is greater than denitrification, allowing a faster growth rate per mole of nitrate compared to denitrification [35]. Thus, as dominant denitrifiers, it would be particularly advantageous for Burkholderiales to evolve higher nitrate turnover when substrate availability is high, allowing more efficient energy production from anaerobic respiration. This might explain the observed Burkholderiales-specific positive selection on Nar nitrate reductase.

### Biological and environmental significance of positive selection

Our inference on the direction of selection relied on codon-wise d_N_/d_S_ ratios (ω). Although this approach has its limitations, discussed below, regions along a gene with strong signals of positive selection are likely to be of biological relevance. This has been clearly demonstrated for protein domains expected to be rapidly evolving, such as those involved in evolutionary arms races. For instance, codons corresponding to protein regions targeted by antibiotics [28], or domains at the interface of host–pathogen interactions [36, 37], tend to form clusters with elevated ω values, indicating positive selection amidst a background of purifying selection (ω < 1). Strong signals of ω > 1 can therefore be useful indicators of adaptive evolution across diverse genetic landscapes and populations.

Despite this, however, we acknowledge the inherent limitations of our approach. d_N_/d_S_ models characterise natural selection based on mutational bias. This approach is limited when synonymous codons drive differential fitness effects caused, for example, by codon usage bias [38]. This leads to potential inflation of ω values, even under purifying selection [38, 39]. However, by analysing rates of positive selection within taxonomic lineages, we could limit the effects of codon-usage biases, mutation rate, and non-neutral synonymous codons on ω, which are inherently linked to taxonomy. Further, by comparing codon-wise ω values for the same gene across different land systems, we could address these potential biases under the assumption that they are consistent across land systems sampled at a global scale. Finally, standardising codon alignment positions and d_N_/d_S_ model parameters ensured uniformity across all land system-specific analyses. This is clearly demonstrated by the fact that most genes analysed had near identical codon-wise ω values across land systems (Supplementary Figs. S2-S3).

Our findings suggest that enzyme kinetics of the Burkholderiales respiratory nitrate reductase is evolving towards increased nitrate turnover. The potential for agricultural practices to alter the evolutionary trajectories of microbial enzymes has important environmental and ecological implications, particularly considering the global scale of our analysis. Croplands account for ∼10% of the Earth’s land surface, with an increasing trend in net primary production due to intensified agricultural land use [40]. Increasing rates of nitrate reduction can have cascading environmental impacts, such as reducing fertiliser-use efficiency and soil health, and exacerbating nitrous oxide emissions – a potent greenhouse gas produced during denitrification. Thus, the overlooked evolutionary consequence of agriculture on microbial proteins and biogeochemistry may have widespread consequences relevant to climate change, food production, ecosystem health, and the success of terrestrial restoration projects. Understanding how agriculture shapes microbial enzyme evolution is therefore critical for predicting feedbacks between land use, climate change, and ecosystem stability.

## Conclusion

Our findings provide the first evidence of anthropogenic selection pressures influencing the evolution of microbial nitrogen cycling processes at a global scale. Specifically, our findings offer novel insights into the potential impact of agriculture on rates of positive selection for respiratory nitrate reductases among Burkholderiales globally. Structural analyses suggest that protein regions involved in enzyme-substrate interactions are undergoing evolutionary change, which could be altering Nar enzyme kinetics, and increasing rates of nitrate turnover in agricultural soils, with important ecological and environmental implications. It therefore may be prudent to incorporate potential evolutionary effects of agrochemicals into regional or planetary boundaries to prevent unmanageable shifts in Earth system health and functioning.

## Methods

### Screening soil metagenomes for nitrogen and phosphorus cycling genes

Metagenomic data were retrieved from the IMG/M database [41]. All assembled soil Illumina-sequenced metagenomes with a GOLD [42] Ecosystem Subtype classification of ‘Agricultural land’, ‘Grasslands’, ‘Temperate forest’, or ‘Tropical forest’ were downloaded on 2023-06-26 (*n* = 2,545 metagenomes; Fig. 1a; Supplementary Table S1). IMG-provided amino acid sequences from each metagenome were screened for nitrogen and phosphorus cycling proteins using a compiled list of KEGG Ortholog (KO) identifiers [43] (Supplementary Table S2). KOs were detected using KofamScan v1.3.0, applying model-specific homology thresholds [44]. Only KOs present in at least 10 metagenomes within each of the four land systems were retained for further analyses. This resulted in 20 nitrogen and 18 phosphorus cycling genes (Supplementary Table S3).

Note, that in the KEGG Orthology database, the catalytic subunits of Nar nitrate reductase (NarG) and Nxr nitrite oxidoreductase (NxrA) are included within the same KO profile (K00370) due to sequence homology. Similarly, the electron transfer subunits of these enzymes, NarH and NxrB, are also included in the one KO profile (K00371). We distinguished protein sequences belonging to each of these subunits based on phylogenetic and protein structural analyses (Supplementary Methods).

### Generating codon-based alignments for ω estimation

Our analysis workflow for estimating ω is illustrated in Fig. 1b. To infer the strength and direction of selection upon that gene, we estimated d_N_/d_S_ (denoted as ω) on a per codon basis. To do this, we first generated a codon-based alignment (Fig. 1b). This involved aligning all protein sequences for a given KO using MUSCLE5 v5.1 [45]. We applied the MUSCLE5 PPP algorithm [parameters: -align] for KOs with less than 2,500 sequences, and the Super5 algorithm [parameters: -super5] for KOs with more than 2,500 sequences. TrimAL v1.4.rev15 [46] was used to trim the protein alignments using the *gappyout* model, and to ‘backtranslate’ these to codon alignments using the corresponding nucleotide sequence for each protein, thus preventing any alignment gaps within a codon [parameters: -fasta -gappyout -backtrans]. This resulted in a single, trimmed, codon-based alignment for each KO. These alignments were then split by land system, resulting in four codon alignments per KO. This approach ensured that the positions of aligned codons were identical across all land systems for a given KO.

### Estimating ω on a per codon basis

We estimated ω on a per codon basis for each gene within each land system using genomegaMap v1.0.1 [28], applying the Bayesian sliding window model. We employed identical parameter settings across all land systems to allow direct comparisons for each KO. We set an exponential prior distribution with a mean of 1.0 for ω, and improper log-uniform prior distributions were set for κ (transition:transversion ratio) and θ (diversity parameter).

GenomegaMap was then run for each codon alignment with 500,000 iterations and a sliding window with mean block length of 30 codons. Output was thinned to retain every 100 iterations, with the first 50,000 iterations discarded as a burn-in.

At each codon, genomegaMap estimates a single ω value from each iteration, resulting in a range of ω values per codon. From this range, we calculated a single ω point estimate (based on the posterior median of all values), a posterior 95% credibility interval, and the posterior probability that ω is greater than one at that codon (Pr(ω > 1)). A codon was classified as being under positive selection if the posterior median of ω exceeded 1, and the corresponding 95% posterior credibility interval did not encompass 1.

### Taxon-specific ω analyses

Genes identified to have a significantly greater proportion of codons under positive selection in a specific land system were re-analysed within taxonomic lineages. For this, the genes were first taxonomically labelled by classifying the whole contig carrying each gene of interest using MMSeqs2 Release 14-7e284 [47, 48] [parameters: taxonomy --tax-lineage 1] against the Genome Taxonomy Database [49-52]. The MMSeqs2 taxonomy workflow assigns contig-level classifications by weighted voting of taxonomic labels assigned to all protein fragments on a contig. For each gene, we separated sequences into all order-level taxa that comprised at least 5% of the sequences for that gene. We then re-calculated codon-wise ω for sequences within each gene-taxon-land system grouping. For the taxon-specific analyses, we employed 1,000,000 iterations of genomegaMap. Duplicate runs were performed for a subset of genes to assess the number of iterations until convergence was observed. This informed the number of iterations to discard as burn-in, which was set to 100,000.

### NarG and NarH protein structural analyses

Experimentally determined structures of NarG and NarH were downloaded from the Protein Data Bank (PDB: https://www.rcsb.org/). To map sites under positive selection to the NarG and NarH crystal structures, the corresponding sequences of each were incorporated into the *narG* and *narH* codon alignments using MAFFT v7.508 [parameters: --add --keeplength -- compactmapout --localpair]. Protein structures were visualised using PyMOL (https://pymol.org/2/). Sites with a posterior probability of positive selection greater than 0.9 (i.e., Pr(ω > 1) > 0.9) were highlighted on the visualised structures.

### Statistical analyses

All statistical analyses were performed using R [53]. For each gene, we tested for significant differences in the proportion of codons under positive selection between land systems through pairwise χ^2^ tests [54]. To account for False Discovery Rate, the resulting p-values were adjusted using the Benjamini-Hochberg correction method [55]. To test for significant differences in NarG amino acid physicochemical properties, we employed non-parametric Wilcoxon rank sum tests with continuity correction [56, 57] as the data significantly deviated from normal distributions (Shapiro-Wilk tests [58], p < 0.001).

## Supporting information

Supplementary Methods, Supplementary Results, Supplementary Figs. S1 to S7

## Data availability

Assembled metagenomic data used in this study are available in the IMG/M database at https://img.jgi.doe.gov/cgi-bin/m/main.cgi.

## Supporting Information

Supplementary Methods

Supplementary Results

Supplementary Figs. S1 to S7

Supplementary Tables S1-S3

## Acknowledgements

TMG would like to thank Mary, Saoirse and Maebh Ghaly for their loving support.

