## Supplementary Methods, Supplementary Results, Supplementary Figs. S1 to S7 for "Agriculture alters protein evolution of nitrogen cycling genes in soil bacteria at a global scale"

**The PDF file (pages S1-S11) includes:**

Supplementary Methods
Supplementary Results
Supplementary Figs. S1 to S7

**Other Supplementary Information for this manuscript include the following:**

Supplementary Tables S1-S3

### Supplementary Methods

*Distinguishing NarGH sequences from NxrAB*

We distinguished NarG protein sequences from NxrA, and NarH from NxrB, based on phylogenetic and protein structural analyses. This first involved obtaining a reference sequence for each subunit that had an experimentally determined function as well as a solved crystal structure. These were NarG (PDB: 1R27_A) and NarH (1R27_B) from *Escherichia coli* [30], and NxrA (7B04_B) and NxrB (7B04_A) from *Kuenenia stuttgartiensis* [59].

Phylogenetic trees (WAG+CAT+Gamma20) were then generated for K00370 metagenomic sequences together with the NarG and NxrA references, and for K00371 metagenomic sequences with the NarH and NxrB references, using FastTree v2.1.11 [60]. From these trees, phylogenetic relatedness between metagenomic sequences and each reference protein was inferred based on cophenetic distances, calculated using the *cophenetic.phylo* function from the R package ape v5.6.2 [61]. Based on cophenetic clustering, six K00370 (NarG/NxrA; Supplementary Fig. S7a) and twelve K00371 (NarH/NxrB; Supplementary Fig. S7d) sequences were selected for AlphaFold2 structural predictions, using ColabFold v1.5.2  [62, 63]. Structural similarity of the predicted structures to each of the reference crystal structures was determined using TM-align [64]. Metagenomic sequences that clustered with each reference sequence, based on phylogenetic and structural homology (Supplementary Fig. S7), was assigned to that gene annotation.

### Supplementary Results

*Distinguishing NarGH sequences from NxrAB*

We found that all predicted K00370 (NarG/NxrA) structures were near identical (TM-score > 0.9) to the reference NarG crystal structure, despite an amino acid sequence identity as low 34% (Supplementary Fig. S7b). Further, we found that, at least for the current dataset, structural homology with either NarG or NxrA increases with phylogenetic relatedness, almost perfectly fitting a quadratic regression ($r^2 = 0.974$; Supplementary Fig. S7c). This suggests that all K00370 sequences in our dataset will likely form near identical structures (TM-score > 0.9) to NarG. We therefore classified all K00370 sequences as NarG nitrate reductases, rather than NxrA nitrite oxidoreductases.

K00371 (NarH/NxrB) metagenomic sequences formed two main clusters based on their cophenetic distances to either NarH or NxrB (Supplementary Fig. S7d), suggesting that these sequences can be distinguished phylogenetically. Structural predictions of K00371 sequences confirmed that these two phylogenetic clusters indeed correspond with structural homology to the crystal structures of either NarH or NxrB ($r^2 = 0.667$; Supplementary Fig. S7d,e). We therefore assigned gene annotations to the K00371 sequences based on the phylogenetic and structural clusters. Two sequences were identified as outliers (Supplementary Fig. S7c,d), and were removed from the dataset prior to $\omega$ calculations.

### Supplementary Figures

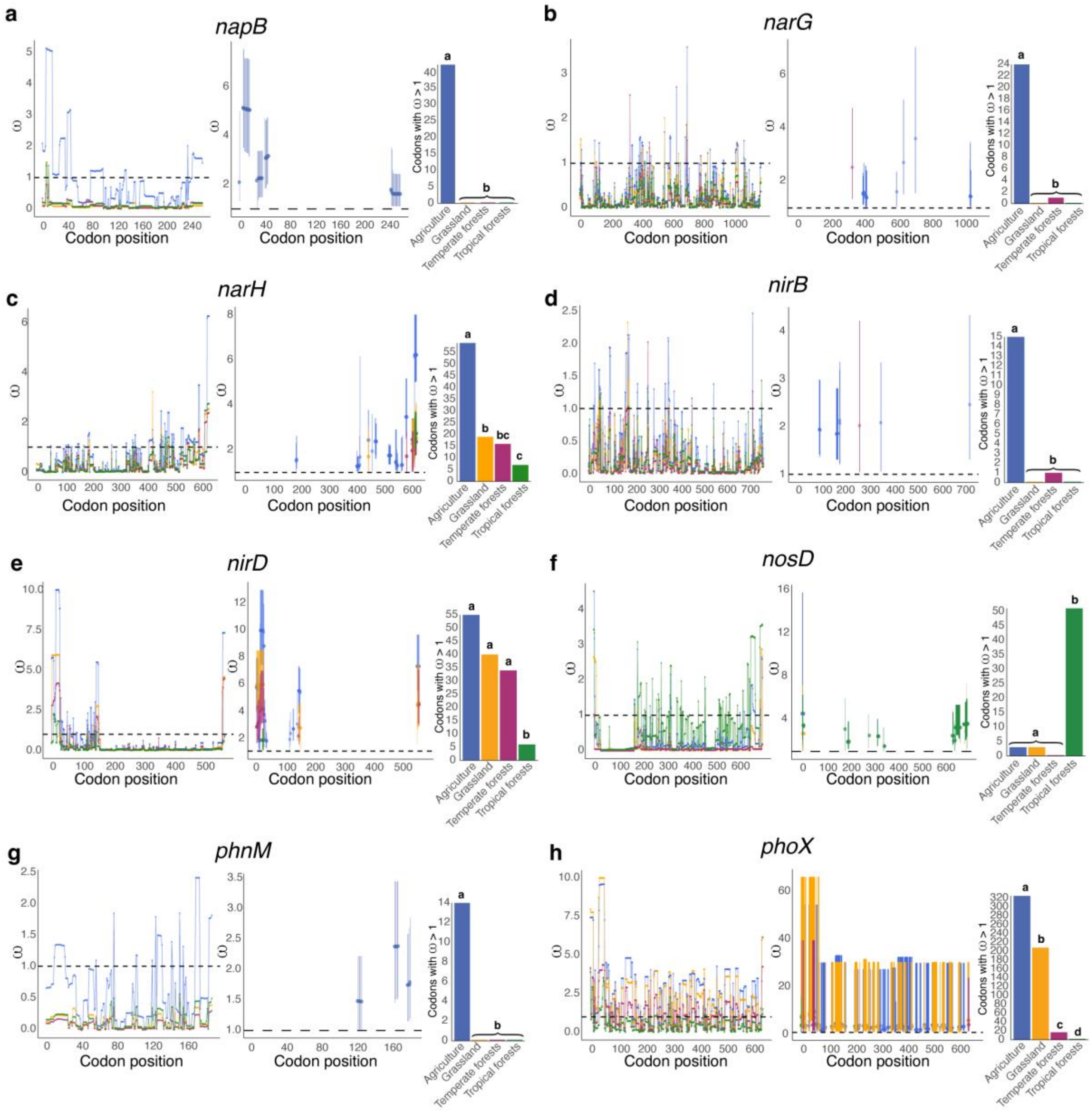


**Supplementary Fig. S1. Genes with significantly different proportions of codons under positive selection between land systems. a-h**, display codons under positive selection for all statistically significant genes highlighted in Fig. 1c. Within each panel: left figure displays posterior median ω values per codon along the gene, coloured by land system; middle figure shows all codons under positive selection, with points representing posterior median ω values, and bars represent their corresponding 95% credibility intervals; right figure shows the number of codons under positive selection for each land system. Statistically significant groupings (lettering above bars) were determined by pairwise $\chi^2$ tests (Benjamini-Hochberg adjusted p-values < 0.01).

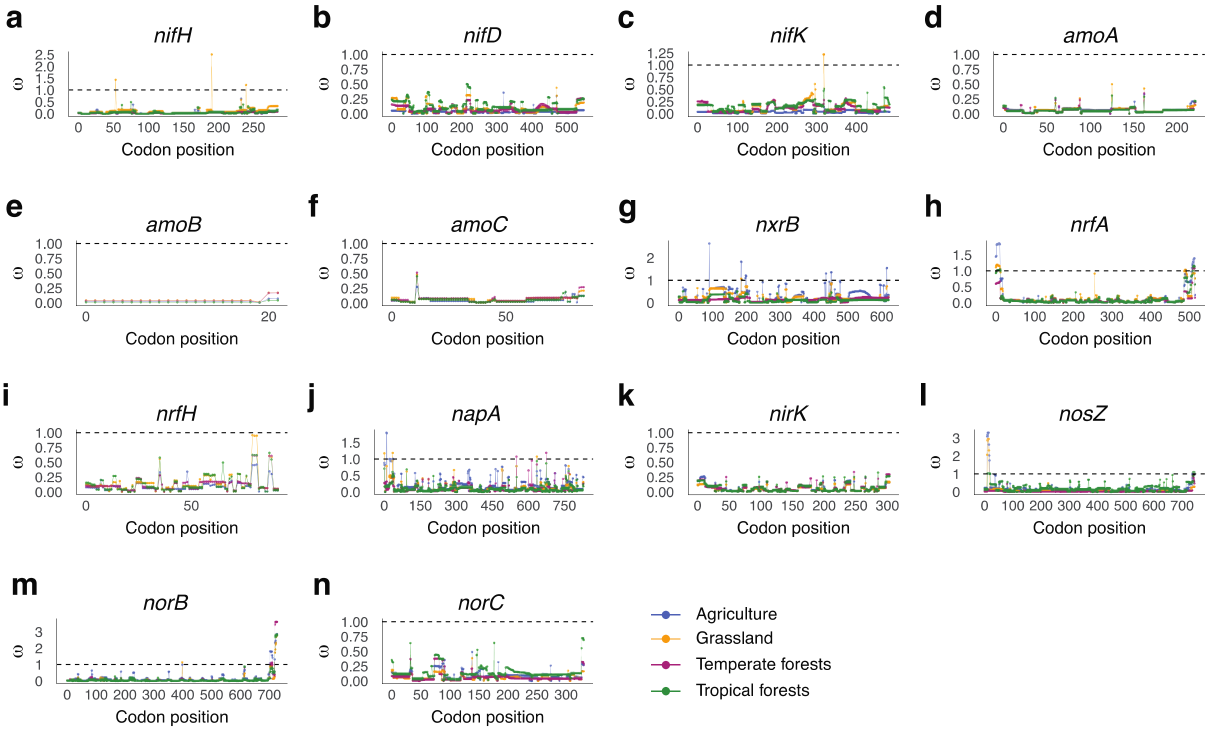


**Supplementary Fig. S2. ω values for non-significant N cycling genes. a-n**, plots display posterior median ω values per codon along each gene, coloured by land system.

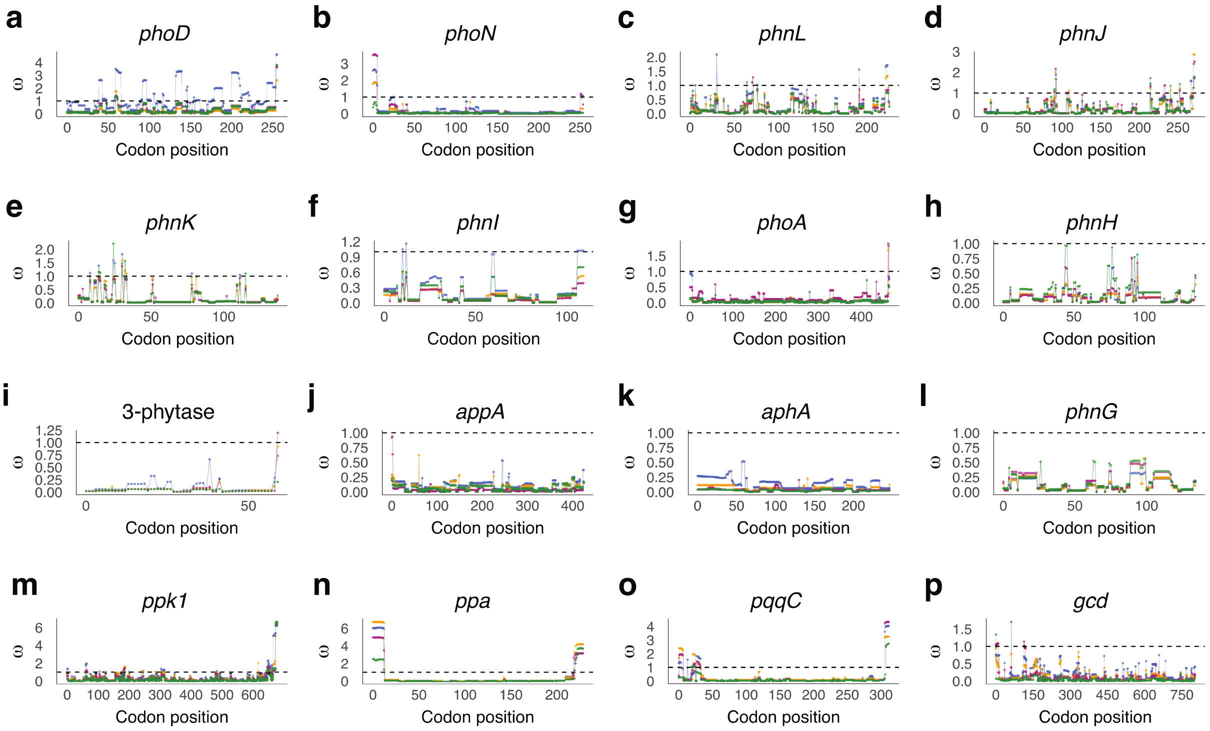


**Supplementary Fig. S3. ω values for non-significant P cycling genes. a-p**, plots display posterior median ω values per codon along each gene, coloured by land system, as per supplementary Fig. S2.

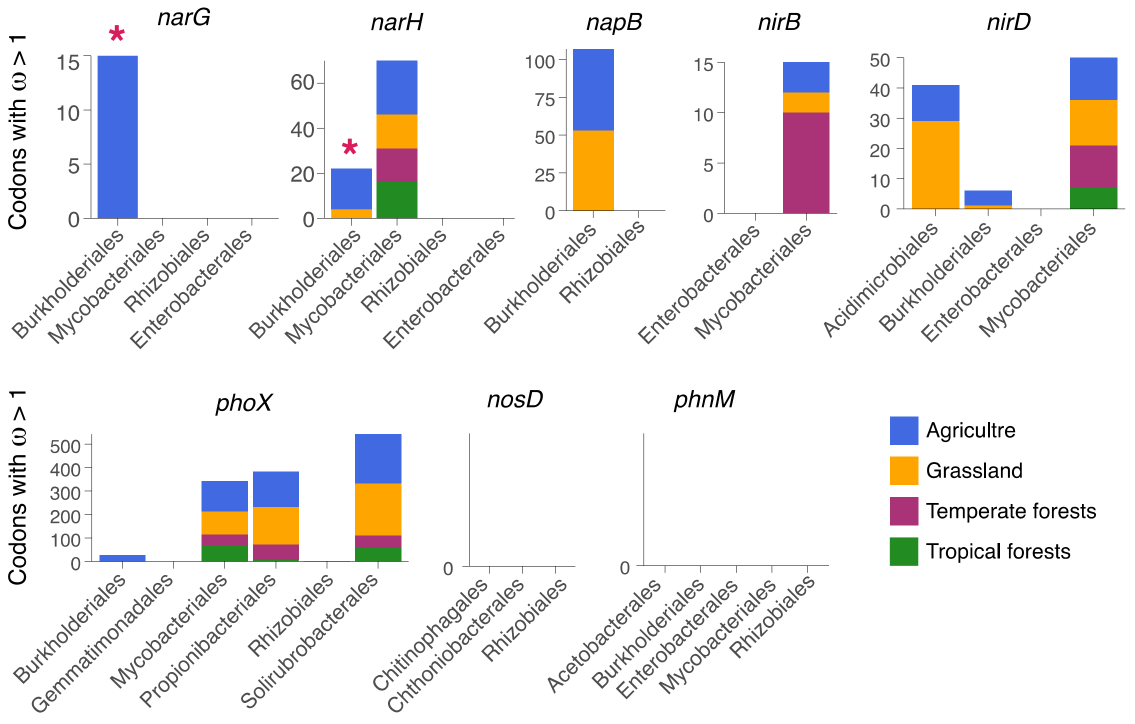


**Supplementary Fig. S4. Land system- and taxon-specific analysis of positive selection.** Stacked bar charts displaying the number of codons under positive selection for each taxon, coloured by land system. Within each taxon, significant difference in the number of codons under positive selection in any one land system are displayed in pink text with asterisks (Pairwise $\chi 2$ tests with Benjamini-Hochberg p-value correction: Burkholderiales narG $p < 0.0008$; and Burkholderiales narH $p < 0.0095$).

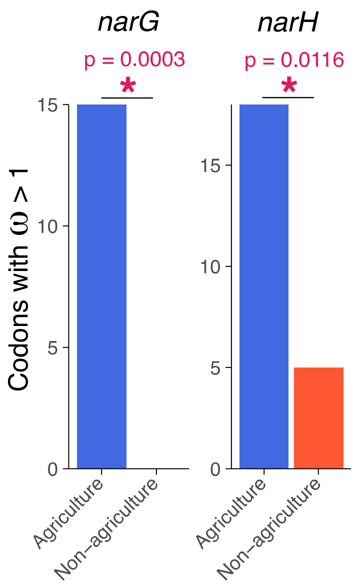


**Supplementary Fig. S5.** The number of Burkholderiales *narG* and *narH* codons under positive selection with agricultural samples versus pooled samples from all natural land systems (non-agriculture). There is a significant increase in the proportions of codons under positive selection in agricultural compared to non-agricultural soils ($\chi^2$ tests, *narG*: p=0.0003, and *narH*: p=0.0116).

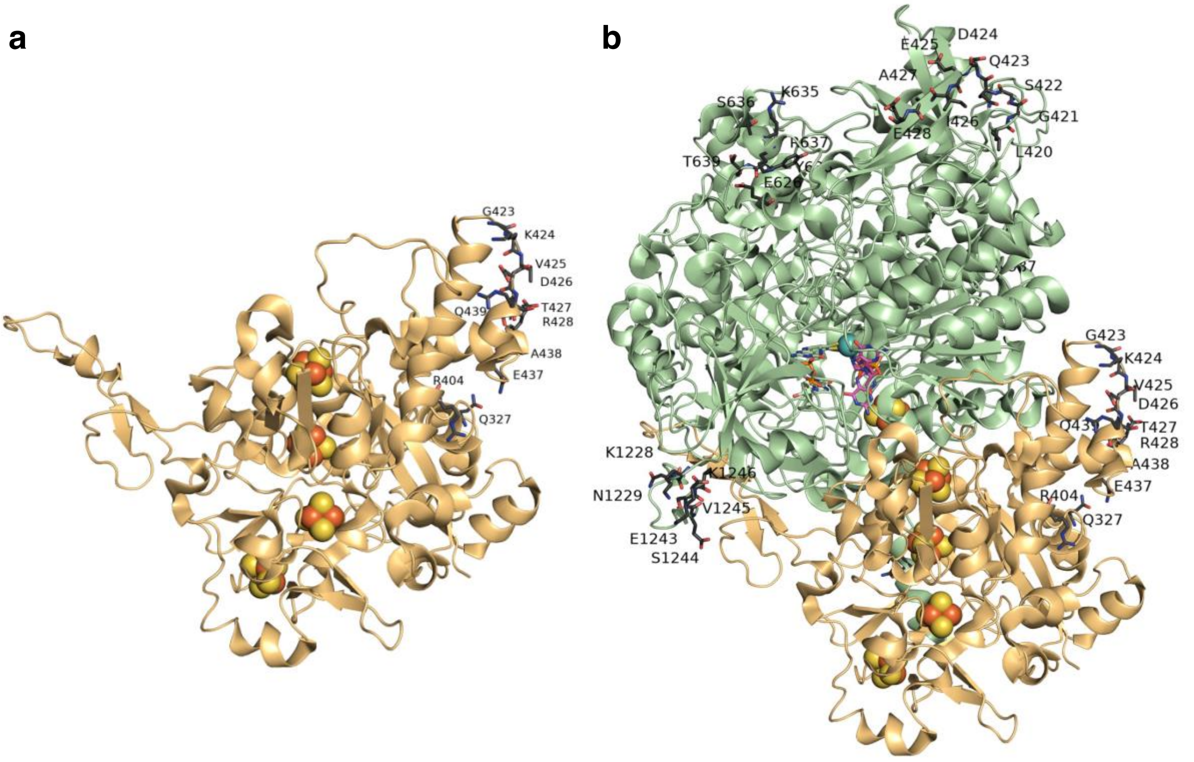


**Supplementary Fig. S6. NarH sites under positive selection in agricultural soils. a**, The NarH crystal structure from *Escherichia coli* (PDB 1R27_B) is depicted as an orange ribbon diagram. Sites under positive selection (i.e., Pr(ω > 1) > 0.9) are shown in black stick representation. The NarH electron-conducting Fe-S clusters are shown as orange and yellow spheres. **b**, The NarG (green ribbon diagram) and NarH (orange ribbon diagram) protein-protein interaction. Neither NarG nor NarH sites under positive selection are in close proximity to the interaction region, or the Fe-S clusters. The functional role of these residues thus remains unclear.

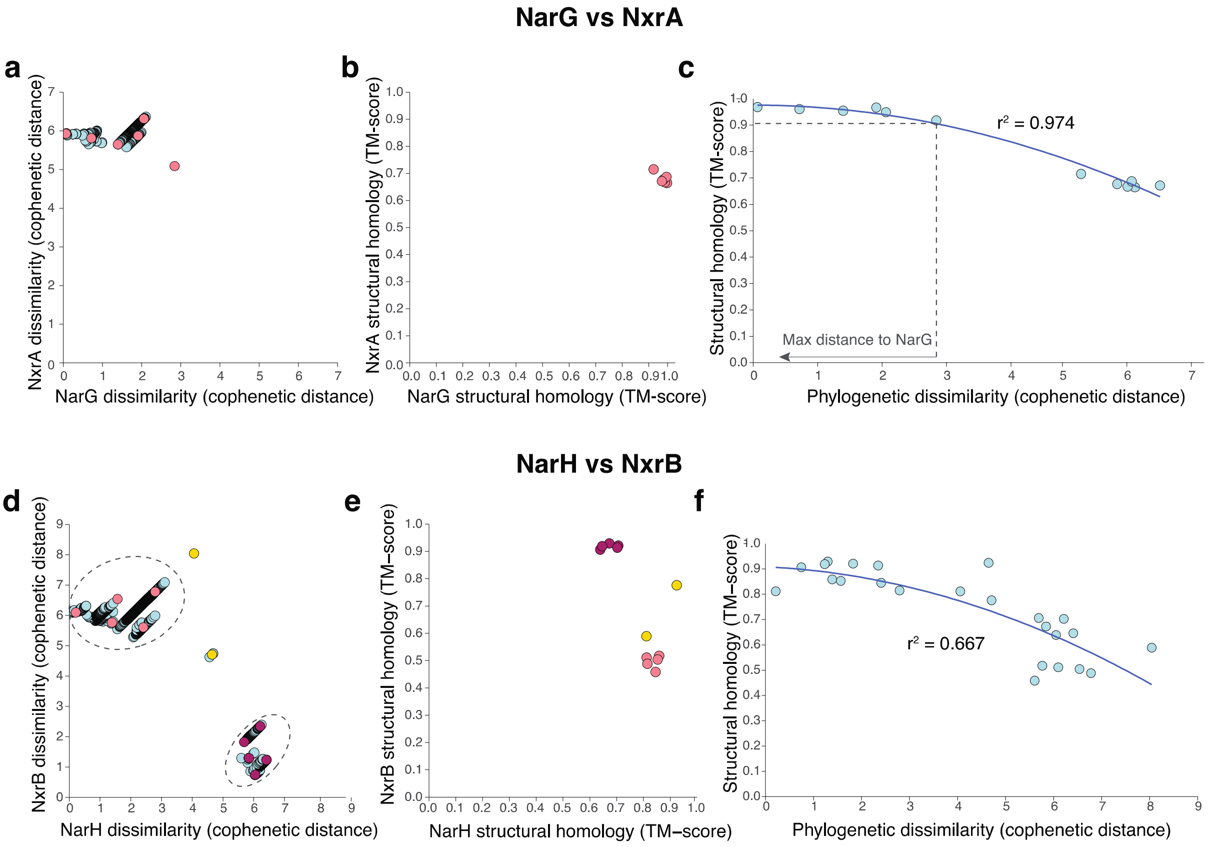


**Supplementary Fig. S7. Distinguishing NarGH sequences from NxrAB.** Phylogenetic and structural data distinguishing **a-c**, NarG from NxrA, and **d-f**, NarH from NxrB. Panels **a** and **d** show the cophenetic distances between metagenomic and reference sequences. Coloured points (not in blue) represent metagenomic sequences selected for structural predictions. **b** and **e** show the structural homology between the predicted metagenomic structures and reference crystal structures. **c** and **f** show the quadratic relationship between cophenetic distance and structural homology between metagenomic and reference proteins.
